# ATHILAfinder: a tool to detect ATHILA LTR retrotransposons in plant genomes

**DOI:** 10.64898/2026.03.20.713144

**Authors:** Elias Primetis, Alexandros Bousios

## Abstract

**Motivation:** The ATHILA lineage of LTR retrotransposons has colonised all branches of the plant tree of life. In *Arabidopsis thaliana* and *A. lyrata*, ATHILA elements have invaded centromeres, influencing the genetic and epigenetic organisation, and driving satellite evolution. To assess the broader significance of ATHILA across plants, a computational pipeline is needed to identify ATHILA elements with high efficiency. Existing tools lack this ability because they are optimised for broad transposon classification at the expense of precise annotation of lower taxonomic levels.

**Results:** We present ATHILAfinder, a pipeline for accurate and large-scale discovery of ATHILA elements. ATHILAfinder uses lineage-specific sequence motifs as seeds and additional filters to build *de novo* intact elements. Homology-based steps rescue intact ATHILA and identify soloLTRs. A detailed identity card includes coordinates, LTR identity, coding capacity, length and other sequence features for every ATHILA. We validate ATHILAfinder in the *A. thaliana* Col-CEN assembly and five additional *Brassicaceae* species, covering four supertribes and ∼30 million years of evolution. ATHILAfinder has very low false positive rates and outperforms widely-used tools like EDTA and the deep-learning-based Inpactor2 software for both recovery and precision of ATHILA. To demonstrate its usefulness, we generate insights into ATHILA dynamics across *Brassicaceae*.

**Outlook:** Few computational pipelines target specific transposon lineages, yet such tools can empower their identification and downstream analyses. Our tailored approach can be adapted to other LTR retrotransposon lineages, offering new ways for high-resolution analysis of transposons.

## Introduction

Transposable elements (TEs) comprise the majority of plant genomic DNA, prompting the development of annotation tools for their analyses (Goerner-Potvin and Bourque 2018). With the advent of next-generation sequencing, these tools facilitated large-scale and automated TE annotation across diverse species. In recent years, long-read technologies have accelerated the production of highly contiguous assemblies, making previously intractable genomes due to their high repetitive content and large size now routinely assembled to high quality (Campos-Dominguez et al. 2025; Niu et al. 2022; Jayakodi et al. 2023). The availability of thousands of assembled genomes has the potential to revolutionise research on genome biology (Shahid and Slotkin 2020), necessitating tools that can identify and analyse TEs in novel ways.

TE annotation is broadly divided into two categories. Homology-based tools like RepeatMasker (Smit A.F.A 2015), CLARI-TE (Daron et al. 2014) and TEannot of the REPET package (Flutre et al. 2011) use libraries of consensus sequences to scan genomes for homologous matches. While effective at detecting degraded or nested TEs, these pipelines are affected by library quality, which requires expert manual curation (Goubert et al. 2022). Even in model species like *Arabidopsis thaliana* and maize, consensus sequences may contain errors or fail to capture native TE diversity (Bousios, Kourmpetis, et al. 2012; Bousios et al. 2017). Additionally, widely-used libraries such as Dfam (Storer et al. 2021) and Repbase (Bao et al. 2015) may underperform in non-model species where TE composition can differ substantially (Baril et al. 2024; Neumann et al. 2019; Platt et al. 2016).

Structure-based methods enable the *de novo* detection of intact TEs at broad taxonomic levels, such as Long Terminal Repeat (LTR) retrotransposons, using conserved structural motifs across eukaryotes (Mhiri et al. 2022). LTR retrotransposons, the most abundant plant TEs, contain two long (∼200-4,000 bp) direct repeats flanking the element, which serve as anchor points for *de novo* pipelines such as LTRharvest (Ellinghaus et al. 2008), LTR_Finder (Xu and Wang 2007), LTR_STRUC (McCarthy and McDonald 2003), and LTRpred (Drost 2020). Post-processing tools like LTRdigest (Steinbiss et al. 2009) and LTRretriever (Ou and Jiang 2018) improve annotation and classify elements into *Ty1/Copia* and *Ty3/Gypsy* superfamilies. Similar principles apply to other TE orders, including Helitrons (Xiong et al. 2014), DNA transposons (Han and Wessler 2010; Su et al. 2019), and Pack-TYPE transposons (Gisby and Catoni 2022). These methods are often combined in pipelines such as EDTA (Ou, Su, Liao, Chougule, Ware, et al. 2019), Repeatmodeler2 (Flynn et al. 2020), and EarlGrey (Baril et al. 2024) to annotate all TEs in a single step. However, these pipelines often produce high false positive and false negative rates, misclassification, and poor boundary annotation (Goubert et al. 2022; Ou, Su, Liao, Chougule, Agda, et al. 2019), which, in turn, affect the performance of homology-based tools that rely on their output.

An alternative approach that can circumvent these issues is developing tools tailored to specific transposon clades at lower taxonomic levels. When these clades are deeply rooted in host phylogeny, these tools can enable insightful cross-species comparisons. Crucially, clades must contain conserved, clade-specific sequence signatures for large-scale detection with high sensitivity and precision. Superfamilies like *Ty1/Copia* and *Ty3/Gypsy* are too diverse for precise annotation, while family-level classification (e.g., ATHILA5, Evade, Opie, Ji) is typically restricted to single or very closely related species, limiting cross-species applicability (Neumann et al. 2019; Ramachandran et al. 2020) (**Figure 1**). In plants, an ideal target lies between the superfamily and family classifications, hereafter called LTR retrotransposon lineages (Neumann et al. 2019). These lineages are phylogenetically distinct, evolutionarily conserved across plants, and possess characteristic coding and non-coding features. A recent survey of 80 hosts revealed 13 *Ty1/Copia* and 12 *Ty3/Gypsy* lineages conserved across the plant kingdom (**Figure 1**) (Neumann et al. 2019). Among them, the SIRE lineage contains multiple conserved non-coding motifs, which enabled the development of the MASiVE algorithm (Bousios, Minga, et al. 2012; Darzentas et al. 2010), uncovering dramatic SIRE expansion in maize (∼20% of genomic DNA) and an evolutionary arms race with epigenetic silencing pathways (Bousios, Kourmpetis, et al. 2012; Bousios et al. 2016).

**Figure 1.**
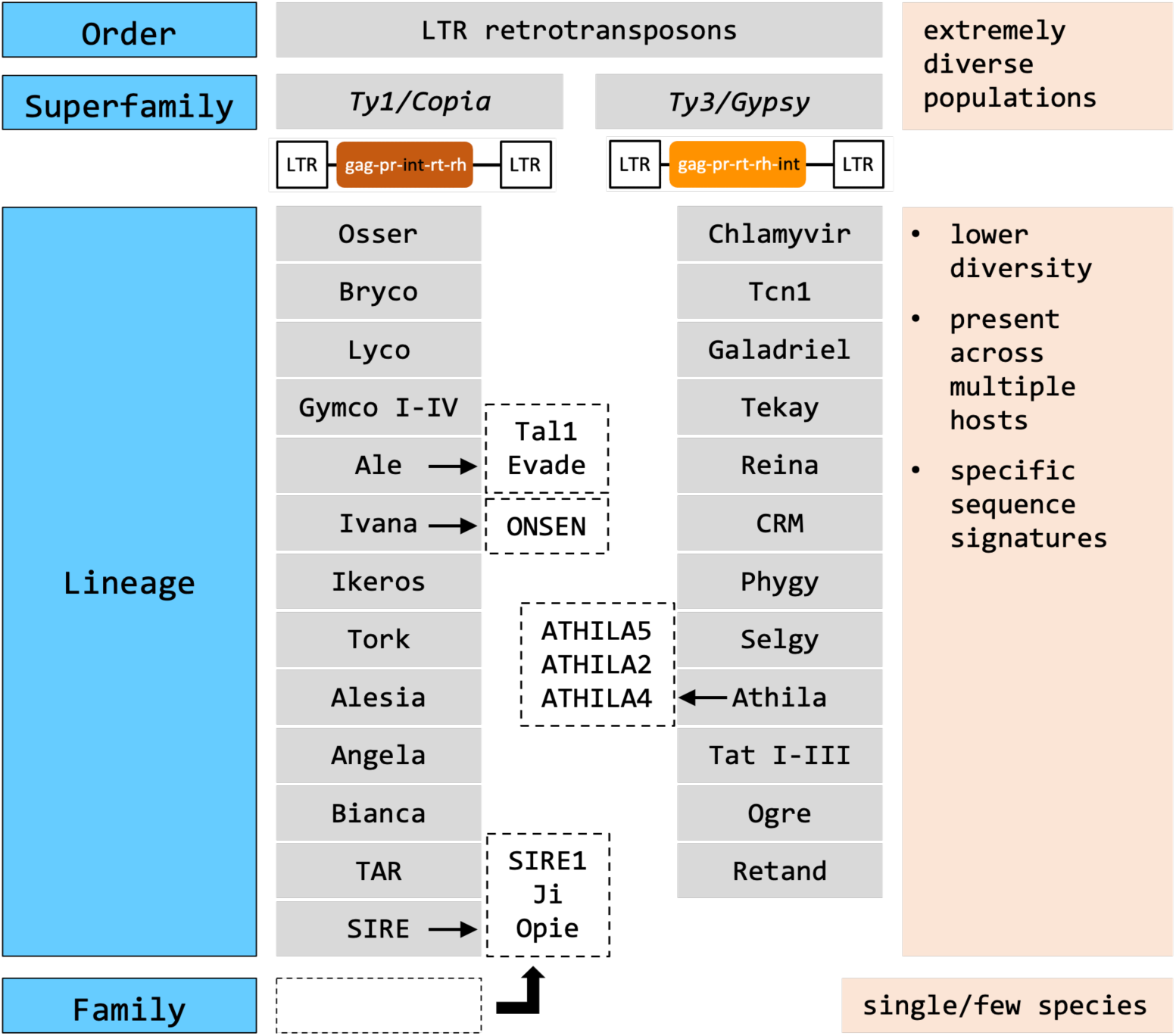
Classification of LTR retrotransposons in plants. *Ty1/Copia* and *Ty3/Gypsy* superfamilies are distinguished by the order of genes in their coding domains. Examples of known families are shown in boxes with dashed lines. pr, protease; int, integrase; rt, reverse transcriptase; rh, RNAseH.

Until recently, few studies have investigated TEs at the lineage level, limiting our understanding of their impact on host genomes. This is changing with tools like TEsorter (Zhang et al. 2022) and DANTE_LTR (Novák et al. 2024), which use hidden Markov model (HMM) profiles to classify intact TEs into lineages. However, ATHILA annotation in the *A. thaliana* Col-CEN assembly required tedious manual curation to rescue elements missed by EDTA and TEsorter (Naish et al. 2021). We therefore developed ATHILAfinder, a dedicated pipeline for the large-scale and accurate discovery of intact ATHILA elements. We describe ATHILAfinder’s design, demonstrate its application in several *Brassicaceae* species, and benchmark its performance against state-of-the-art annotators like EDTA (Ou, Su, Liao, Chougule, Agda, et al. 2019) and Inpactor2 (Orozco-Arias et al. 2023). We also discuss the broader potential of lineage-specific pipelines to improve transposon annotation and facilitate functional and evolutionary analyses.

## Methods

### Identification of *ATHILA*-specific sequence motifs

At the core of ATHILAfinder are sequence motifs that we detected in the LTR-internal junctions, which are specific to ATHILA and conserved across hosts. These motifs serve as seeds during genome scans, with downstream steps generating high-quality intact elements (**Figure 2**). We found these conserved junctions, by initially generating a population of intact ATHILA elements with EDTA (--anno 1) and TEsorter (-db rexdb-plant -nolib on *Ty3/Gypsy* elements) in six *Brassicaceae* species spanning 135 to 367 megabase pairs (Mbp) and four supertribes over ∼30 million years of evolution: *A. thaliana* (Naish et al. 2021), *A. lyrata* (Wlodzimierz et al. 2023), and *Erysimum cheiranthoides* (Züst et al. 2020) from *Camelinodae*, *Brassica nigra* (Paritosh et al. 2021) from *Brassicodae*, *Megadenia pygmaea* (Yang et al. 2021) from *Heliophilodae*, and *Draba nivalis* (Nowak et al. 2021) from *Arabodae* (**Table S1**). This process identified 903 ATHILA (49-340 elements per species) (**Table 1**), which were merged in a single dataset with manually curated ATHILA from Col-CEN (Naish et al. 2021). We then performed multiple alignments of the first and last 2.5 kbp of these elements (MAFFT v7.453, --retree 2 --maxiterate 1000) (Katoh and Standley 2013) to ensure that the LTR-internal junctions were captured.

**Figure 2.**
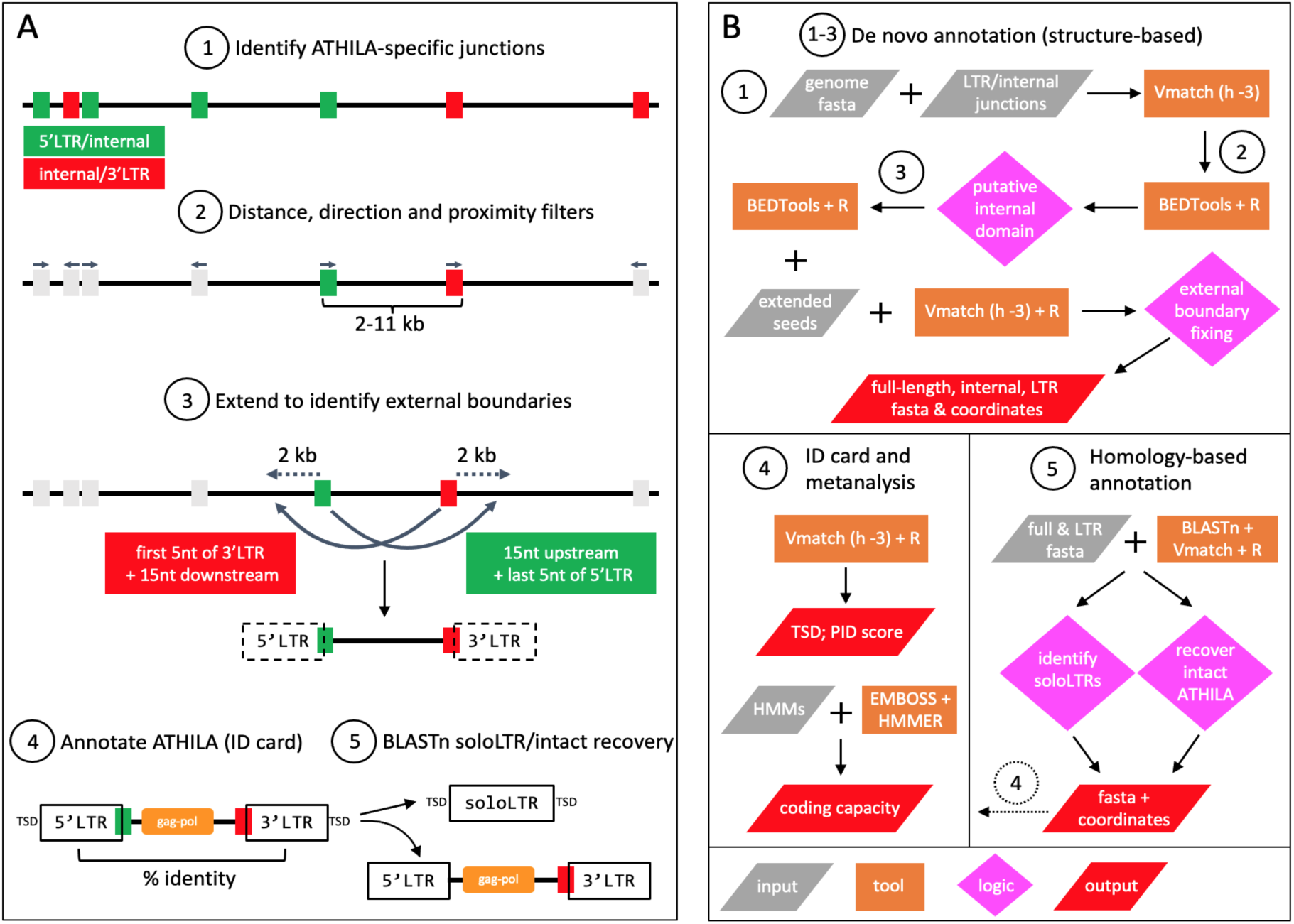
The ATHILAfinder pipeline. The left panel (**A**) depicts the workflow at the genome level, the right panel (**B**) summarises the computational tasks. ATHILAfinder consists of three main modules: i) *de novo* structural identification of intact ATHILA elements (steps 1-3 in both panels). The genome fasta file and the sequence seeds of the LTR-internal junctions are provided as input; ii) Meta-analysis of the intact elements to generate positional, sequence, coding, and LTR identity information (step 4); iii) Homology-based recovery of intact ATHILA and identification of soloLTRs (step 5), using as input intact ATHILA and their LTRs from the *de novo* step. Search and filtering parameters can be adjusted by the user, as indicated in Table 2.

**Table 1.** ATHILA annotation by EDTA/TEsorter, Inpactor2 and ATHILAfinder across six

| TE pipeline | TE types | *Camelinodae* |  |  | *Brassicodae* | *Heliophilodae* | *Arabodae* |
| --- | --- | --- | --- | --- | --- | --- | --- |
|  |  | *A. thaliana* | *A. lyrata* | *E. cheiranthoides* | *B. nigra* | *M. pygmaea* | *D. nivalis* |
| EDTA | Ty1/Copia | 108 | 699 | 556 | 814 | 269 | 364 |
|  | Ty3/Gypsy | 96 | 829 | 447 | 371 | 222 | 539 |
|  | LTR retrotransposons | 85 | 200 | 272 | 214 | 97 | 165 |
|  | TEsorter-ATHILA | 49 | 340 | 241 | 77 | 80 | 116 |
| Inpactor2 | Ty1/Copia | 62 | 251 | 261 | 401 | 126 | 299 |
|  | Ty3/Gypsy | 529 | 558 | 246 | 302 | 236 | 483 |
|  | ATHILA | 25 | 166 | 108 | 38 | 48 | 31 |
| ATHILAFinder | ATHILA (total) | 144 | 539 | 497 | 375 | 392 | 375 |
|  | *de novo* | 140 | 489 | 437 | 359 | 279 | 346 |
|  | homology | 4 | 50 | 60 | 16 | 113 | 29 |
|  | soloLTR | 151 | 206 | 333 | 474 | 2136 | 824 |
|  | *de novo* (TSD %) | 70.0 | 85.5 | 80.3 | 64.9 | 63.8 | 57.8 |
|  | homology (TSD %) | 25.0 | 54.0 | 51.7 | 43.8 | 48.7 | 44.8 |
|  | soloLTR (TSD %) | 71.5 | 75.2 | 75.1 | 61.6 | 50.0 | 60.1 |

**Table 2:**
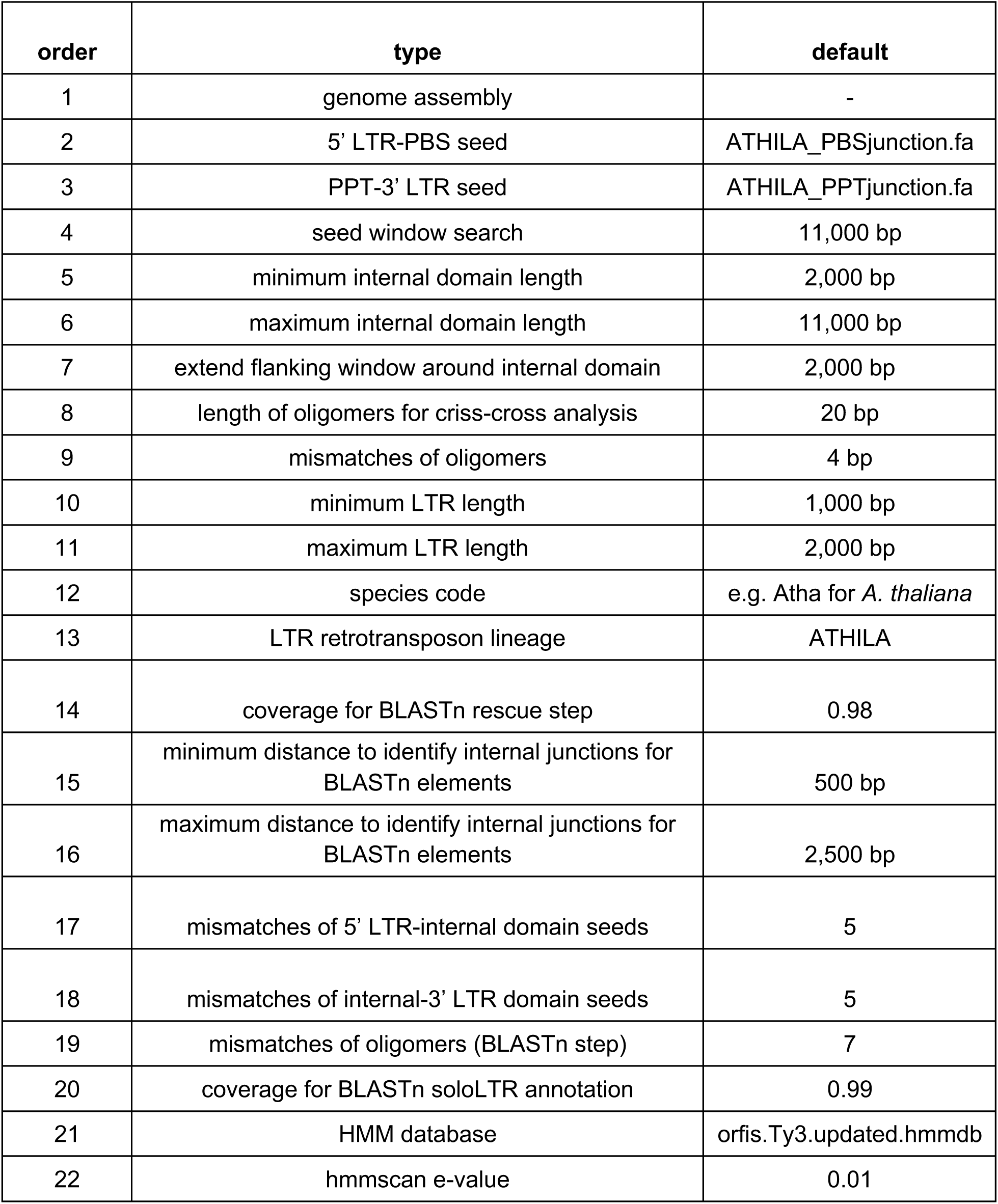
ATHILAfinder parameters that can be adjusted by the user.

### Brassicaceae species

Based on this approach, we were able to identify and design seeds for both junctions. The 5’ LTR-internal junction contained a conserved LTR terminus and the primer binding site (PBS) for tRNA-Asn and/or tRNA-Asp that share the same TGGCGCCGTTGCC 3’ sequence (**Figure 3A**). The LTR terminus and PBS were separated by a short intervening region, forming the attachment site that is important in template switching during reverse transcription (**Figure 3A, S1**) (Bousios et al. 2010; Hughes 2015). Although the intervening region varied across species, it was based on an AT-rich 3-6bp motif. The most common 5’ LTR-internal junction contained the TATCA LTR terminal pentamer followed by the ATT intervening trinucleotide in 504 elements (56%) across five species (**Figure 3A, S1**). We designed six 21bp seeds that always included the 5’ LTR terminal pentamer to aid precise external boundary identification (**Figure 2**). The internal-3’ LTR junction was highly conserved across nearly all ATHILA (97%). Two polymorphic positions occurred in the sequence upstream of the polypurine tract (PPT) (**Figure 3B**): C/T at position 1 with species-specific variation, and position 6 with T dominant (94-100%) in *A. thaliana* and *A. lyrata* versus C dominant (70-94%) in the other species. We designed three 20bp seeds accounting for these variants, always including the 3’ LTR upstream pentamer.

**Figure 3:**
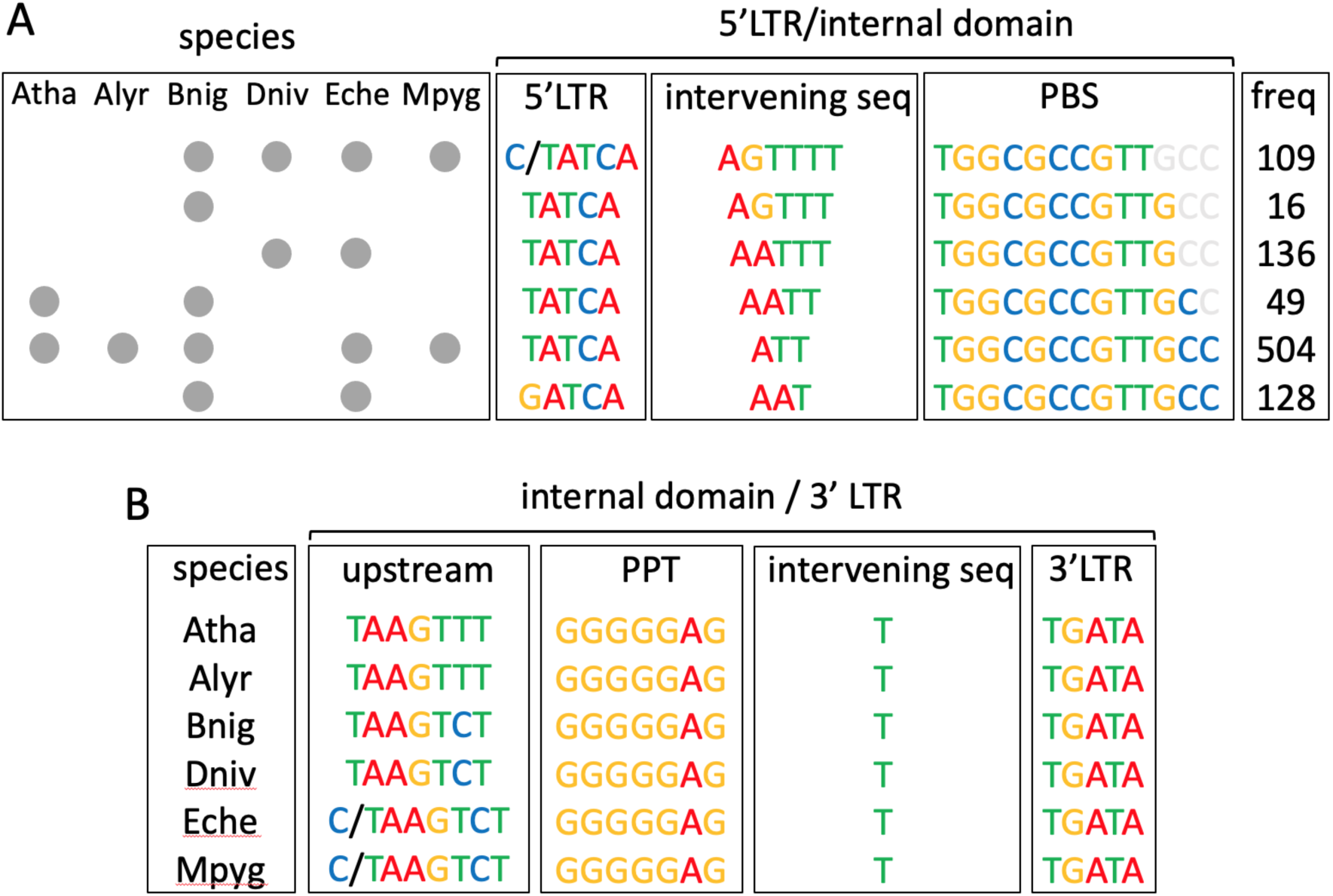
Conservation of LTR-internal junctions in ATHILA. Sequence structure of the 5’ LTR-internal (**A**) and internal-3’ LTR (**B**) junctions. Only sequences that appear in >10% of the ATHILA dataset are shown. In A, species that contain a seed are highlighted with a grey dot. Nucleotides of the PBS in light grey are not part of the seeds. Atha, *A. thaliana*; Alyr, *A. lyrata*; Bnig, *B. nigra*; Dniv, *D. nivalis*; Eche, *E. cheiranthoides*; Mpyg, *M. pygmaea*.

Importantly, we verified seed specificity to ATHILA. Using Vmatch (http://www.vmatch.de/), we searched 16,548 EDTA-derived intact TEs from all superfamilies not classified as ATHILA by TEsorter. With 4 mismatches (-h 4), only 454 TEs (2.7%) generated hits with the seeds, of which 182 contained both LTR-internal junctions. Some of these TEs likely contain ATHILA fragments or are misclassified intact ATHILA that lack coding sequence, which is common in *A. thaliana* and *A. lyrata* (Wlodzimierz et al. 2023). This analysis confirms that the LTR-internal junctions are conserved and specific to ATHILA within *Brassicaceae*.

### Building *de novo* intact ATHILA elements

To detect intact ATHILA *de novo*, ATHILAfinder first parses the genome fasta file (treat_fasta.py) to standardise the format, and then uses Vmatch (h -3) to identify the seeds (step 1 in **Figure 2**). We identified 912 (*A. thaliana*) to 3,539 (*B. nigra*) 5’ LTR-PBS and PPT-3’ LTR seeds combined (**Figure 4**), which were filtered to retain pairs within 2-11 kbp in the same orientation and correct order, generating putative internal domains (step 2 in **Figure 2**). Loci containing additional seeds internally or in flanking 1 kbp windows were rejected, as they may represent nested structures or proximal ATHILA with incomplete LTRs. This produced 3,002 internal domains ranging from 200 (*A. thaliana*) to 603 (*D. nivalis)* (**Figure 4**).

**Figure 4:**
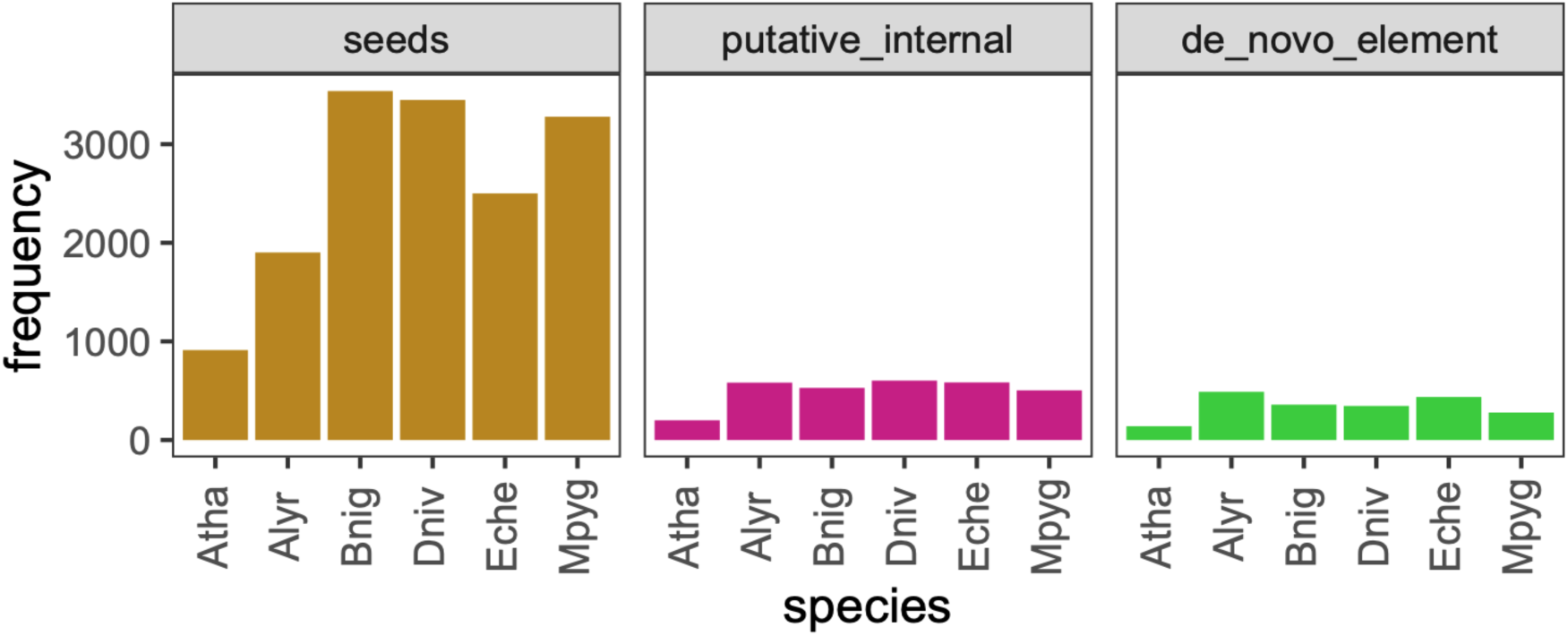
***De novo* structural annotation workflow of ATHILAfinder.** The left panel shows the total number of non-redundant Vmatch hits of the 5’ LTR-internal and internal-3’ LTR seeds in the six *Brassicaceae* species (corresponds to step 1 in Figure 2). In the middle panel, these hits are parsed by various filters to generate putative internal domains of 2-11 kbp length (step 2 in Figure 2). In the right panel, additional steps identify the external borders of elements and produce the final set of structurally intact ATHILA (step 3 in Figure 2). Atha, *A. thaliana*; Alyr, *A. lyrata*; Bnig, *B. nigra*; Dniv, *D. nivalis*; Eche, *E. cheiranthoides*; Mpyg, *M. pygmaea*.

ATHILAfinder next identifies the external borders of putative internal domains to verify the presence of flanking LTRs (step 3 in **Figure 2**). Precise TE boundary identification is challenging (Goubert et al. 2022), but ATHILAfinder achieves nucleotide resolution by leveraging the fact that the seeds contain the first and last 5 bp of LTRs (**Figure 3**). We generate two 20mers per putative internal domain, composed of the LTR pentamer (from the seed) and adjacent outward-facing 15 nucleotides. These fragments, precisely located at LTR extremities, are used in ‘criss-cross’ mode to detect external boundaries (step 3 in **Figure 2**). Using Vmatch (-h 4), we search windows located 1-2 kbp from the putative internal domain, based on the typical ATHILA LTR length of >1 kbp (Naish et al. 2021; Wlodzimierz et al. 2023; Neumann et al. 2019), selecting the hit with the fewest mismatches. This completes the *de novo* identification of full-length ATHILA, totalling 2,050 intact elements across six species, ranging from 140 in *A. thaliana* to 489 in *A. lyrata* (**Figure 4**, **Table 1**).

### Homology-based rescue and soloLTR identification

As structural annotation may miss elements (e.g. due to deletions spanning the seeds), we applied a BLASTn-based homology approach to rescue intact ATHILA (step 5 in **Figure 2**). After masking genomes for already annotated intact elements, we used them as queries in BLASTn runs, requiring 98% coverage that corresponds to a ∼9.8 kbp alignment for a 10 kbp element. Retrieved loci were processed similarly to *de novo* elements. We searched for the LTR-internal junctions using the ATHILA seeds (Vmatch, h -5) and applied the 20mer criss-cross step (Vmatch, h -7) for external boundary identification. Overall, we retrieved 272 ATHILA (**Table 1**), most (∼80%) with 100% query coverage, indicating high-quality annotation (**Figure S2**).

The annotation of external and internal boundaries with nucleotide resolution also facilitates the identification of soloLTRs, products of unequal homologous recombination between the LTRs of a single element or of highly similar proximal elements (Cossu et al. 2017). This type of recombination is common, given the abundance of soloLTRs across hosts, and drives TE deletion in genomes (Cossu et al. 2017; Kent et al. 2017). Identification of soloLTRs, however, is largely *ad hoc* with no dedicated tool available during whole-genome TE annotation. ATHILAfinder uses intact ATHILA LTRs as queries (step 5 in **Figure 2**), masks intact elements, and runs BLASTn with 99% coverage threshold, corresponding to a 1.485 kbp alignment for a 1.5 kbp LTR. We extract the first and last 50bp of the target loci (i.e. beginning and end of the LTR) plus 50bp of flanking regions, forming two 100bp fragments. Using the LTR-internal seeds (Vmatch, -h 5), we scan these fragments for internal domain evidence, and classify them as soloLTRs when no hits are identified. This generated 4,124 soloLTRs across six species (**Table 1**).

### Meta-analysis of *ATHILA* and output files

ATHILAfinder generates detailed data and metrics for every ATHILA element and soloLTR in a main tab-delimited output file (step 4 in **Figure 2**). Information includes target site duplications (TSDs), reported by allowing one mismatch in flanking pentamers. We explore coding capacity by scanning ORFs ≥300bp with HMMER (v3.3.2, --E 0.001 --domE 0.001) (http://hmmer.org/), using Hidden Markov Models (HMMs) from Pfam (Mistry et al. 2021) (**Table S2**). These HMMs describe core domains of the *gag* gene and *pol* polyprotein, plus an ATHILA-specific domain downstream of *pol* (Slotkin 2010). ATHILAfinder also calculates the Percentage Identity (PID) Score of LTR pairs using the *pid* function (Biostrings package, R). All data appear in the main output file together with genomic coordinates, chromosome number, orientation, and full/LTR length. Intact ATHILA and soloLTRs receive unique IDs (including a species code) that are preserved across output files. Fasta files are generated for all ATHILA features, and information on intermediate steps is available in various formats (fasta, bed, Vmatch).

### Parameters of ATHILAfinder

Several parameters can be adjusted by the user in the command line (**Table 2**). These include (1) input genome fasta, (2,3) 5’ LTR-PBS and PPT-3’ LTR seeds fasta, (4) window around each seed where matching pairs are formed; (5,6) minimum/maximum length of the putative internal domain. The default is 2-11 kbp, but testing is advised for species that may contain ATHILA with longer internal domains; (7) window around the putative internal domain to filter out loci with flanking additional seeds; (8,9) oligomer length and mismatches during the criss-cross step. We tested various combinations (15-25 bp and -h 2 to 4) and found that 20mers and 4 mismatches provide the best balance of specificity and sensitivity. If additional seeds are designed, we advise that the first and last LTR pentamers are always included, as they guide precise boundary identification during the criss-cross step; (10,11) minimum/maximum LTR length, defaulting to 1-2 kbp based on recent work in *A. thaliana* and other plant species (Naish et al. 2021; Wlodzimierz et al. 2023; Neumann et al. 2019) ; (12) species code across output files and (13) LTR retrotransposon lineage name. Avoid uncommon characters as they can trigger bugs; (14) coverage threshold for the BLASTn rescue step (default 0.98). Lower thresholds increase recovery at the expense of quality; (15,16) minimum/maximum distances from the beginning/end of the element to identify internal junctions in the BLASTn rescue step (500 and 2,500 bp default); (17,18) mismatches allowed in the 5’ and 3’ junction seeds in the BLASTn rescue step; (19) mismatches allowed in the oligomers for the external boundary annotation of BLASTn-rescued elements; (20) coverage threshold for soloLTR identification (default 0.99). Lower thresholds increase recovery but also the false discovery rate; (21) HMM database, adjustable if required; (22) e-value used during hmmscan.

## Results

### *ATHILA* elements are identified with very low false positive rates

We tested the performance of ATHILAfinder by estimating the false positive rate. We built phylogenetic trees based on the *gag* and *reverse transcriptase* core domains of *Ty3/Gypsy* LTR retrotransposons in a dataset that included i) ATHILAfinder intact elements from all six species (**Table 1**), ii) *Ty3*/*Gypsy* elements identified by EDTA and assigned a lineage by TEsorter in all six species (**Table 1**), and iii) 151 known ATHILA from Col-CEN (Naish et al. 2021). EDTA elements and known ATHILA provided the framework for placing ATHILAfinder elements. A single ATHILA branch formed in both trees, separated from other *Ty3/Gypsy* lineages such as CRM, Galadriel, Reina, Retand and Tekay (**Figure 5A,B**). Apart from seven and five sequences in the gag and reverse transcriptase trees respectively, all ATHILAfinder elements are co-located in the ATHILA branch together with ATHILA from Col-CEN and EDTA.

**Figure 5:**
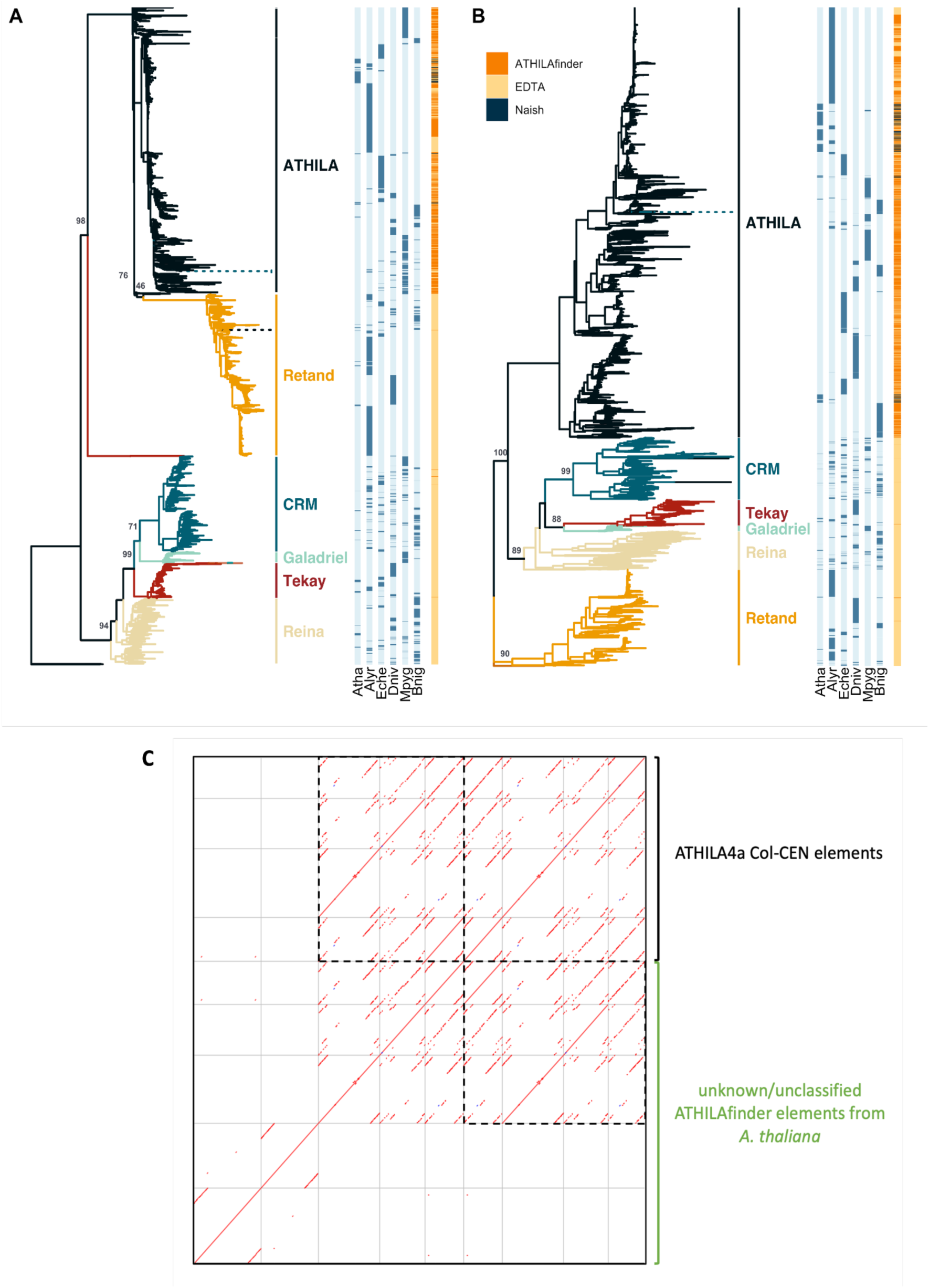
Phylogenetic analysis of *Ty3/Gypsy* lineages in six *Brassicaceae* species. (**A,B**) The phylogeny was based on the reverse transcriptase (PF00078) (A) and gag (PF03732) (B) core domains. Elements of all datasets were run through TEsorter to retrieve the coding domains: 1,885 with gag and 647 with reverse transcriptase for ATHILAfinder; 2,391 and 1,557 for EDTA; 142 and 32 for known ATHILA (Naish et al. 2021). The low number of ATHILA with the reverse transcriptase gene is due to the prevalence of non-autonomous intact elements with a ∼3 kbp internal deletion in the coding region (Wlodzimierz et al. 2023). The amino acid sequences were aligned with MAFFT (--retree 2 --maxiterate 1000) (Katoh and Standley 2013), and phylogenetic trees were produced with FastTree (-nt) (Price et al. 2009). Bootstrap support of key nodes is shown. Dotted lines correspond to long tips that were capped to aid visualisation. The trees were rooted using the maize Huck *Ty3/Gypsy* element. (**C**) Dot plot full-length sequence comparison between five ATHILAfinder elements from *A. thaliana* that were unclassified/not parsed by TEsorter and four known ATHILA4a elements. Window size is 20bp. Atha, *A. thaliana*; Alyr, *A. lyrata*; Bnig, *B. nigra*; Dniv, *D. nivalis*; Eche, *E. cheiranthoides*; Mpyg, *M. pygmaea*

We also ran the ATHILAfinder dataset through TEsorter, and the vast majority were classified as ATHILA (1,966/2,322, 85%). Among the remaining elements, most were Ty3/unknown (143, 6%) or not parsed by TEsorter (135, 6%), both outcomes resulting from a complete lack of HMM detection. These elements likely belong to non-autonomous ATHILA families lacking coding capacity, such as ATHILA4a and some ATHILA0 elements in *A. thaliana* (Wlodzimierz et al. 2023). We retrieved five elements from *A. thaliana* classified as Ty3/unknown or not parsed, and dot plot analysis showed that three belonged to ATHILA4a (**Figure 5C**). Overall, the tree topology and high sequence similarity with known non-autonomous ATHILA families strongly support ATHILAfinder’s specificity.

### Benchmarking ATHILAfinder with general annotation pipelines

ATHILAfinder identified 2,322 intact elements in the six *Brassicaceae* species, ranging from 539 ATHILA in *A. lyrata* and 497 in *E. cheiranthoides* to 144 in *A. thaliana* (**Table 1**). The majority were identified *de novo* (2,050, 88%). Across species, ATHILAfinder identified substantially more ATHILA than the state-of-the-art annotator pipelines EDTA/TEsorter (903) and Inpactor2 (416, with parameters -m 17000 -n 2000) (**Table 1**), with Inpactor2 being the first general annotator automatically classifying LTR retrotransposons into lineages (Orozco-Arias et al. 2023). Given ATHILAfinder’s very low false positive rate (**Figure 5**), we intersected the coordinates of the three datasets. Two-thirds of ATHILAfinder elements did not overlap with EDTA/TEsorter or Inpactor2, but, *vice versa*, most EDTA/TEsorter (82%) and Inpactor2 elements (79%) did (**Figure 6a**). We explored if the 215 elements from EDTA/TEsorter and Inpactor2 not intersecting with ATHILAfinder represent false negatives in our pipeline. Indeed, ATHILAfinder did not detect one or both PBS and PPT seeds for 153 elements (29 no PPT, 22 no PBS, 102 no PPT/PBS), possibly due to gaps spanning LTR-internal junctions or small indels within the seeds. This was not the case, however, for the remaining 62 elements that did not satisfy downstream filtering criteria, for example, when additional seeds were found within length windows or when resolving external boundaries (**Figure 2**). Assuming that there are 2,475 (2,322+153) high-quality intact elements across all tools, we calculate a low ATHILAfinder false negative rate of ∼6%.

**Figure 6:**
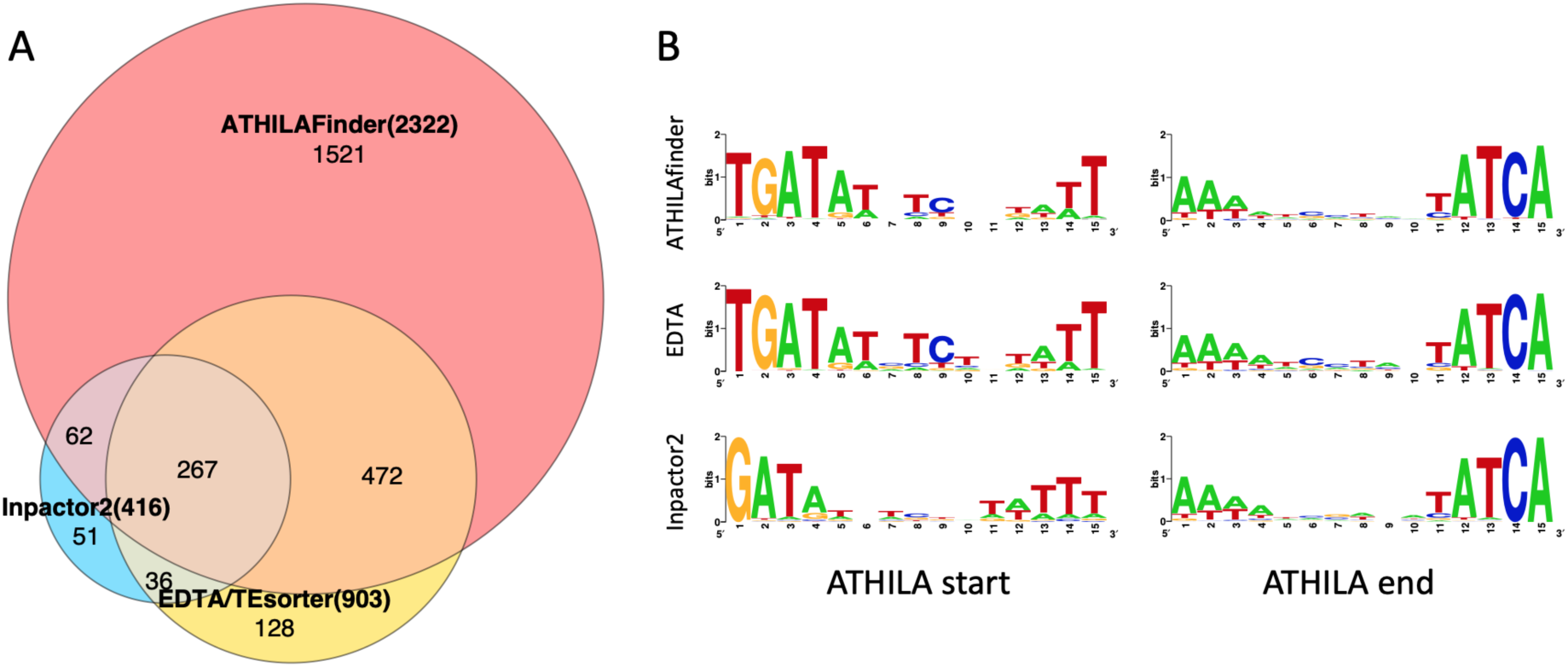
**Comparison between ATHILAfinder, EDTA and Inpactor2 annotation**. (**A**) Intersection of coordinates of intact elements identified by the three pipelines. (**B**) Sequence logos of the upstream (left panel) and downstream (right panel) 15mers of intact elements.

Terminal sequence analysis also showed that ATHILAfinder precisely resolved external and internal LTR boundaries, starting and finishing with universal TG and CA dinucleotides with conserved positions extending inwards (**Figure 6b**). EDTA captured identical termini to ATHILAfinder, while Inpactor2 had lower specificity. We detected TSDs for 72% of intact ATHILA identified *de novo* and 66% of soloLTRs across all species (**Table 1**). The proportion was lower for BLASTn-rescued intact elements (49%), attributed to inefficient delineation of the LTR boundaries in some cases. Users can adjust coverage thresholds for BLASTn searches to balance between quality and quantity (**Table 2**).

### Insights into the dynamics of ATHILA across *Brassicaceae*

The *Brassicaceae* family comprises >4,000 species with nearly 100 fully-assembled species currently available (https://www.plabipd.de/). It was recently proposed as the first plant model clade, with numerous groups studying this family worldwide (Hendriks et al. 2023; Mabry et al. 2024). Within this context, applying ATHILAfinder across six species from four supertribes spanning ∼30 million years provides novel insights into the dynamics of ATHILA within this plant family. Our phylogenetic analysis supports the rapid radiation of ATHILA alongside the evolution of their host species (**Figure 5A,B**). This is corroborated by high LTR identity levels that point towards recent activity in these genomes (**Figure 7A**), and multiple peaks of their full-length sequence profile that suggest the evolution of diverse families (**Figure 7B**). High LTR identity was prominent in the *Camelinodae* supertribe (*A. thaliana*, *A. lyrata*, and *E. cheiranthoides*), and, especially, in *A. lyrata*, which had 80 elements with 100% LTR sequence identity. We note that 29 and 65 of these elements were not identified by EDTA and Inpactor2, respectively.

**Figure 7:**
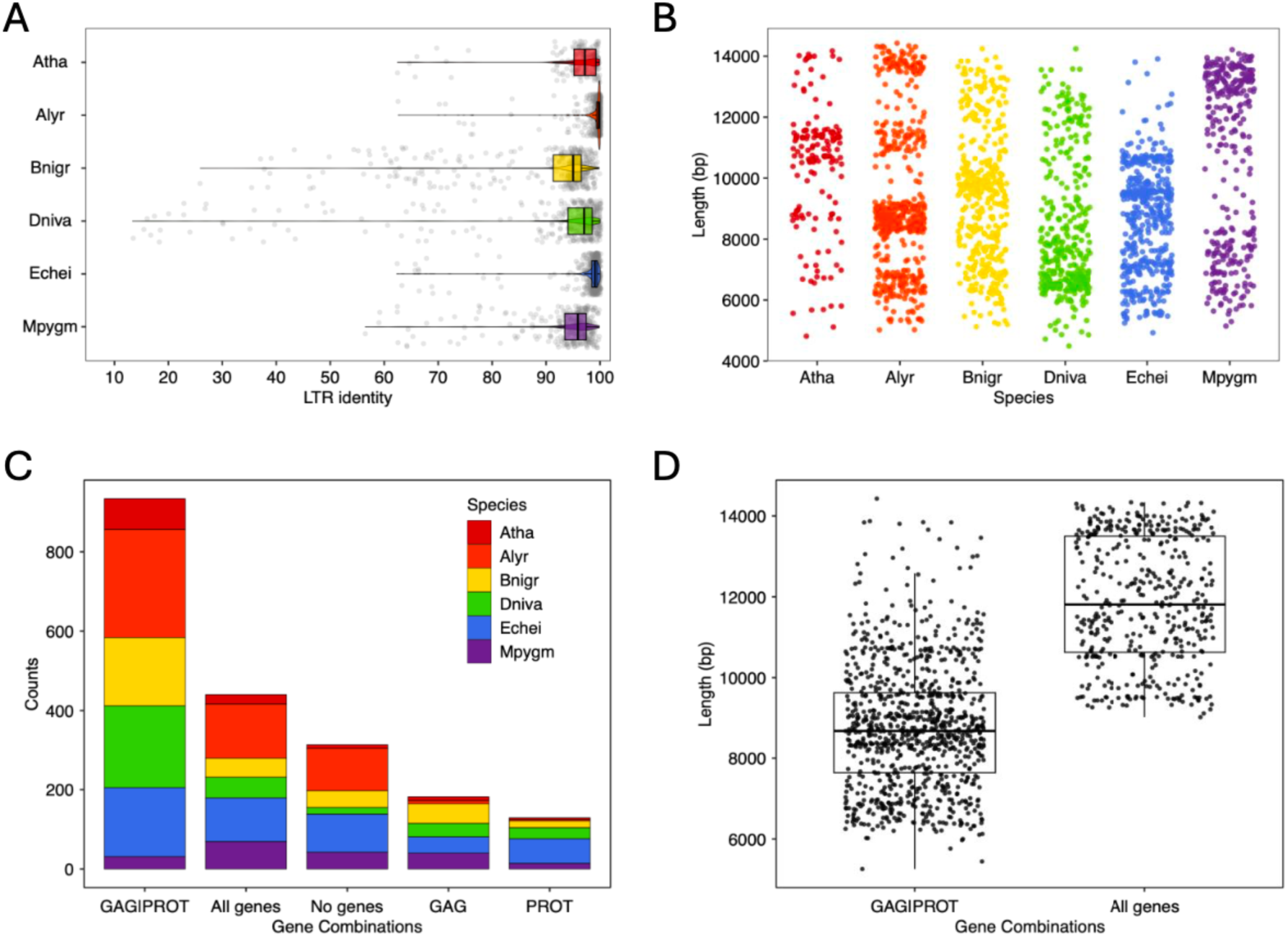
**Characteristics of intact ATHILA in six *Brassicaceae* species.** (**A**) Box plots show the LTR identity values for intact ATHILA. (**B**) Scatter plots show the length of intact ATHILA. (**C**) Bar plots show the five most abundant gene combinations of intact ATHILA based on HMM core domain profiles. (**D**) Box plots show the length distribution of autonomous and non-autonomous intact ATHILA, based on the presence of all five and only the gag-protease HMMs, respectively. All data were retrieved from the identity card of each ATHILA in the main output file of ATHILAfinder. Atha, *A. thaliana*; Alyr, *A. lyrata*; Bnig, *B. nigra*; Dniv, *D. nivalis*; Eche, *E. cheiranthoides*; Mpyg, *M. pygmaea*.

In *A. thaliana*, abundant non-autonomous intact ATHILA contain a large ∼3 kbp deletion that includes the reverse transcriptase, RNaseH and integrase genes (Wlodzimierz et al. 2023). These elements belong to multiple families with little sequence homology, consistent with a conserved recombination hotspot within the *pol* polyprotein, and have been recently active in native populations, including within the centromeres of the ANGE-B-10 accession in France (Wlodzimierz et al. 2023). To test if the same non-autonomous derivative extends beyond *A. thaliana*, we used ATHILAfinder HMM searches (**Table S2**) to classify elements by their *gag* and *pol* genes. Many contained all five genes (440/2,322, 19%), corresponding to autonomous ATHILA, but this was most abundant only in *M. pygmaea* (**Figure 7C**). ATHILA with only gag and protease genes were the most abundant in all other species (934, 40%). The difference in average length between 440 autonomous (11,886 bp) versus 934 gag-protease-only (8,784 bp) elements is 3,102 bp, strongly indicating that this specific non-autonomous type is common across *Brassicaceae* (**Figure 7D**). Around a quarter of elements from all species (673/2,322) have the ATHILA-specific ORF downstream of *gag-pol* (Bousios et al. 2025). The ATHILA ORF currently has no assigned function, which, together with its relatively low frequency, suggests that it is not an essential or highly conserved feature of the lineage. Taken together, these preliminary data show that multiple ATHILA families with complex phylogenetic relationships are currently active across *Brassicaceae*.

## Discussion

Computational tools for TE annotation typically prioritise breadth over depth, identifying TEs at broad taxonomic levels such as superfamilies while sacrificing precision at finer scales. This trade-off limits their utility for high-quality annotation of specific TE lineages across multiple species. ATHILAfinder addresses this gap by identifying ATHILA *Ty3/Gypsy* elements using conserved, lineage-specific sequence motifs as primary seeds. Applied across six *Brassicaceae* species spanning ∼30 million years of evolution, ATHILAfinder demonstrated very low false positive rates (<1% based on the phylogenetic analysis), substantially outperformed general annotators (2.6-fold and 5.6-fold more elements than EDTA/TEsorter and Inpactor2), and enabled first insights into ATHILA dynamics. This includes the discovery that non-autonomous ATHILA derivatives with a characteristic ∼3kb deletion in the coding region are widespread across Brassicaceae. Future work expanding this analysis across additional species and LTR retrotransposon lineages will help determine whether this deletion-based non-autonomy represents a common evolutionary pattern.

ATHILAfinder does have limitations. Elements with degraded LTR-internal junctions may be missed by the seed-based approach. Additionally, the pipeline is optimised for ATHILA within *Brassicaceae*. Extending to other plant taxa may require identifying and updating the conserved motifs within those species. The principles of ATHILAfinder, however, can be applied to other LTR retrotransposon lineages if they exhibit similar conservation patterns. Pipelines designed for detecting particular TE types are currently scarce, including MASiVE for SIRE *Ty1/Copia* elements (Darzentas et al. 2010) and CAULIFINDER for caulimovirid endogenous viral elements in plant genomes (Vassilieff et al. 2022). As sequencing efforts continue to generate high-quality chromosome-level assemblies across the tree of life, such targeted annotation tools can become increasingly valuable for understanding the diverse roles of specific TE lineages in genome function and evolution.

## Data availability

ATHILAfinder is freely available at https://github.com/eliasprim/ATHILAFinder/. The installation of Vmatch, BEDTools, BLAST, EMBOSS, and HMMER algorithms and R, Python and Awk programming languages is required. Data annotation files from running ATHILAfinder in the six *Brassicaceae* species are available in https://doi.org/10.5281/zenodo.19132278.

## Acknowledgements

We acknowledge grant support from Royal Society awards UF160222, RF/ERE/221032, URF/R/221024, RGF/R1/180006, RGF/EA/201030, and RF/ERE/210069 to AB.

## Contributions

AB designed the study and the ATHILAfinder pipeline. EP developed the pipeline. AB and EP wrote the manuscript.

## Supplementary Tables

**Table S1:** *Brassicaceae* species used in ATHILAfinder

| Species | Supertribe | Bioproject | Genome size |
| --- | --- | --- | --- |
| *Arabidopsis thaliana*<br>Col-CEN | Camelinodae | PRJEB46164 | 135 Mbp |
| *Arabidopsis lyrata*<br>NT1 | Camelinodae | PRJEB50329 | 200 Mbp |
| *Brassica nigra* | Brassicodae | PRJNA324621 | 367 Mbp |
| *Draba nivalis* | Arabodae | PRJNA657155 | 276 Mbp |
| *Ersimum*<br>*cheiranthoides* | Camelinodae | PRJNA563696 | 172 Mbp |
| *Megadenia pygmaea* | Heliophilodae | PRJNA637465 | 205 Mbp |

**Table S2:** Hidden Markov Models used in the meta-analysis of ATHILA elements.

| Protein description | Pfam accession code |
| --- | --- |
| Retrotransposon gag protein | PF03732 |
| Gag-polyprotein putative aspartyl protease | PF13975 |
| Aspartyl protease | PF09668 |
| Aspartyl protease | PF13650 |
| Retroviral aspartyl protease | PF00077 |
| Reverse transcriptase (RNA-dependent DNA polymerase) | PF00078 |
| RNAase H-like domain found in reverse transcriptase | PF17917 |
| RNAase H-like domain found in reverse transcriptase | PF17919 |
| Integrase zinc binding domain | PF17921 |
| Integrase core domain | PF00665 |
| Integrase core domain | PF13683 |
| *ATHILA ORF-1 family* | PF03078 |

## Supplementary Figures

**Figure S1:**
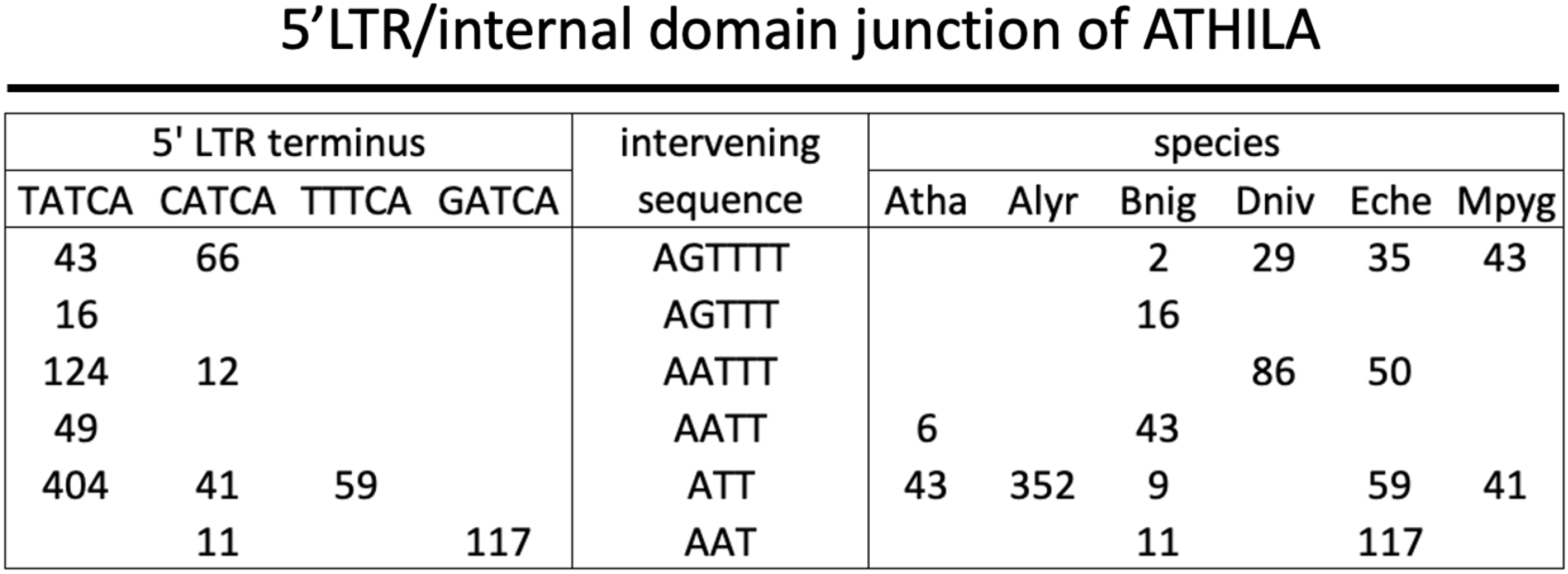
Organization of the 5’ LTR-internal domain junction. Numbers indicate the ATHILA elements within a species with a given combination of the 5’ LTR terminus and the intervening sequence. The PBS is conserved and not shown. 942 out of 1,007 ATHILA (94%) contained an identifiable high-quality junction and were therefore included in this Figure.

**Figure S2:**
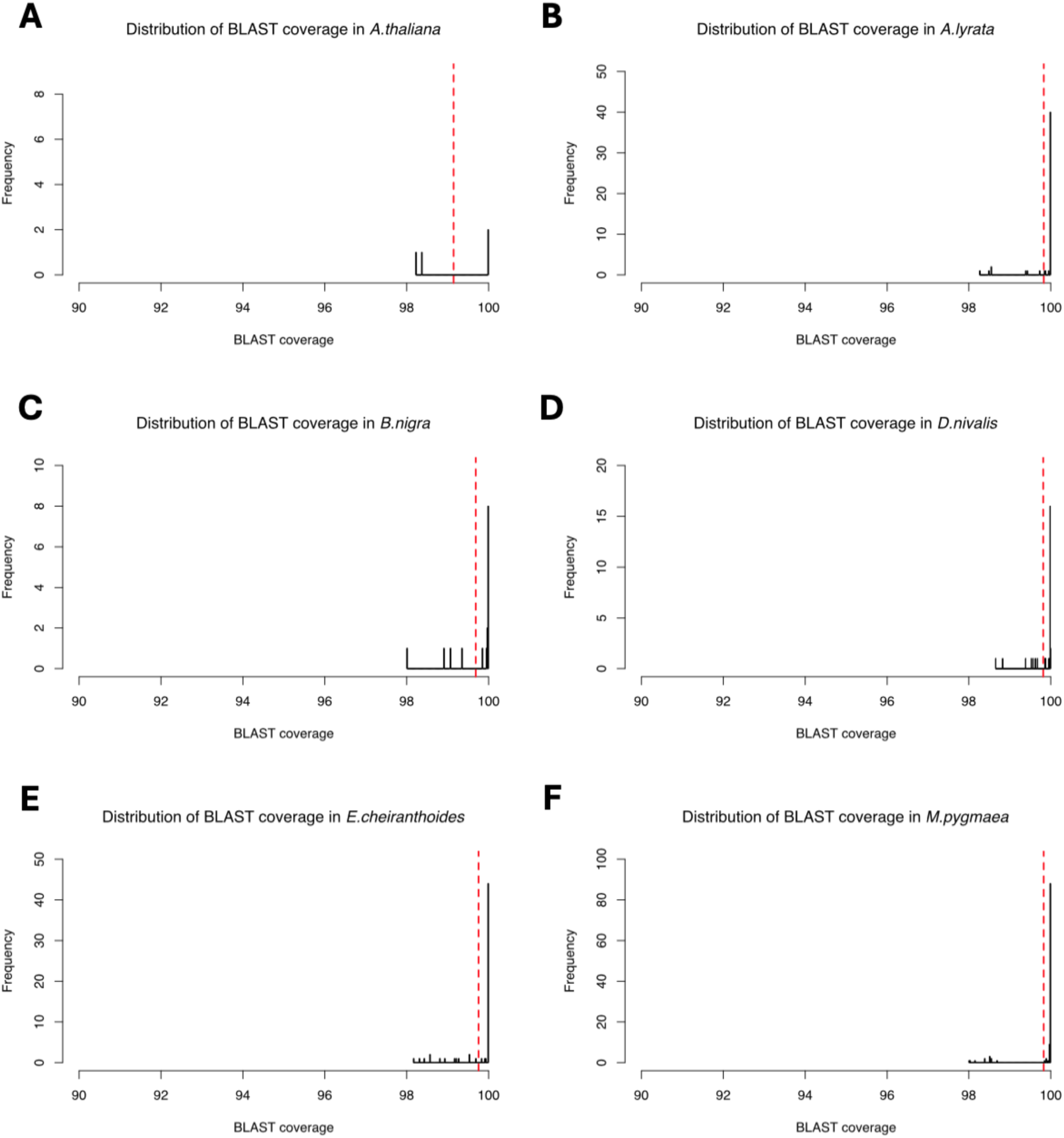
Distribution of BLAST coverage in the six *Brassicaceae* species during the BLASTn-based homology identification. Red dotted lines indicate the mean value.

